# m^6^A Regulates Liver Metabolic Disorders and Hepatogenous Diabetes

**DOI:** 10.1101/2020.01.21.912600

**Authors:** Yuhuan Li, Qingyang Zhang, Guanshen Cui, Fang Zhao, Xin Tian, Bao-Fa Sun, Ying Yang, Wei Li

**Author notes:** Corresponding authors. (Li W), (Yang Y). Equal contribution (Yuhuan Li, Qingyang Zhang).

## Abstract

*N*^6^-methyladenosine (m^6^A) RNA methylation is one of the most abundant modifications on mRNAs and plays an important role in various biological processes. The formation of m^6^A is catalysed by a methyltransferase complex containing a key factor methyltransferase-like 3 (Mettl3). However, the functions of Mettl3 and m^6^A modification in liver lipid and glucose metabolism remain unclear. Here, we show that both *Mettl3* expression and m^6^A level increased in the liver of mice with High Fat Diet (HFD)-induced metabolic disorders, and overexpression of *Mettl3* aggravated HFD-induced liver metabolic disorders and insulin resistance. Hepatocyte-specific knockout of *Mettl3* significantly alleviated HFD-induced metabolic disorders by slowing weight gain, reducing lipid accumulation and improving insulin sensitivity. Mechanistically, Mettl3 depletion-mediated m^6^A loss causes extended RNA half-lives of metabolism-related genes, consequently protects mice against HFD-induced metabolic syndrome. Our findings reveal a critical role of Mettl3-mediated m^6^A in HFD-induced metabolic disorders and hepatogenous diabetes.

## Introduction

*N*^6^-methyladenosine (m^6^A), the most prevalent mRNA modification in eukaryotes [1], is catalyzed by a methyltransferase complex including methyltransferase-like 3 (Mettl3), methyltransferase-like 14 (Mettl14), Wilms’ tumor 1-associating protein (Wtap), among which Mettl3 functions as the catalytic subunit [2, 3]. m^6^A methylation can be reversed by at least two eraser enzymes, fat-mass and obesity-associated protein (Fto) and α-ketoglutarate-dependent dioxygenase alkB homolog 5 (Alkbh5) [4, 5], and is mainly recognized by YTH domain-containing family ‘reader’ proteins [6–10]. As the most abundant and reversible modification on mRNAs, m^6^A has been proved to play key roles in all fundamental aspects of mRNA metabolism such as RNA stability [6] and RNA splicing [8], as well as mRNA translation efficiency [7, 9–11]. Many essential biological processes are known to be regulated by m^6^A, including cell fate determination [12, 13], embryonic development [13–15] and tumorigenesis [16].

Liver plays a central role in the regulation of lipid and glucose metabolism, and is the major site of fatty acid disposal, the main source of endogenous glucose production, and the primary site of insulin degradation [17]. Unhealthy diet habits can lead to liver metabolic disorders, such as Nonalcoholic Fatty Liver Disease (NAFLD), followed by whole-body insulin resistance [17]. Some studies have revealed that m^6^A modulation of mRNA expression is involved in obesity [18] and liver metabolism [19, 20], and plays an important role in the maintenance and progression of liver diseases [21–23]. For instance, a significant increase in FTO mRNA and protein levels has been found in the liver of human NAFLD patients [24]. Elevated levels of FTO mRNA and protein can also be found in a NAFLD rat, which is involved in oxidative stress and lipid deposition [25]. Knockdown of *Mettl3* or *Ythdf2 in vitro* increases the stability and expression of *peroxisome proliferator activator receptor* α (*Ppar*) mRNA, resulting in reduced accumulation of lipids [20]. A recent study showed Mettl3 inhibits hepatic insulin sensitivity via m^6^A located in *Fatty acid synthase (Fasn)* mRNA and promotes fatty acid metabolism [26]. All these studies indicate the important roles of m^6^A in liver metabolic diseases. However, how Mettl3-mediated m^6^A methylation affects liver metabolism and the underlying pathways and mechanisms are still not fully elucidated.

In the present work, we demonstrated that the m^6^A methyltransferase Mettl3 and m^6^A level were consistently up-regulated in the liver from mice after feeding High Fat Diet (HFD). Adeno-associated virus (AAV) mediated liver-specific overexpression of *Mettl3* aggravates liver metabolic disorders and insulin resistance. In turn, we specifically inactivated Mettl3 in the mouse liver using the *Alb*-Cre mediated *Mettl3* conditional knockout (cKO) model and confirmed *Mettl3* ablation protects mice against HFD-induced liver metabolic disorders and insulin resistance, Furthermore, mechanism analysis suggested that *Mettl3* deletion alters expression pattern of liver lipid and glucose metabolism genes, particularly extends mRNA stability of an important regulator of liver metabolism--*Lpin1*. Together, these findings reveal the critical roles for Mettl3 mediated m^6^A modification in HFD-induced liver metabolic disorders and hepatogenous diabetes, supporting that m^6^A could be used as a potential therapeutic and diagnostic target for hepatic diseases.

## Results

### *Mettl3* expression and m^6^A level increased in HFD mice

To explore the potential role of m^6^A in regulation of lipid and glucose metabolism in HFD-induced obese mice, we first measured the relative mRNA levels of m^6^A “writers”, “erasers”, and “readers” in mouse livers after HFD (60 kcal% fat diet) for 20 weeks, including *Mettl3*, *Mettl14*, *Wtap*, *Fto*, *Alkbh5*, *Ythdf1*, *Ythdf2*, *Ythdf3*, *Ythdc1*, *and Ythdc2*. The expression of m^6^A methyltransferases significantly increased in HFD mouse livers, while there was no difference in demethylases or m^6^A binding proteins **(Figure 1A)**. Given Mettl3 is the key ‘writer’ of m^6^A modification [2, 3], we further confirmed the significantly increased protein level of Mettl3 by western blotting and immunohistochemistry assays **(Figure 1B-D)**.

**Figure 1.**
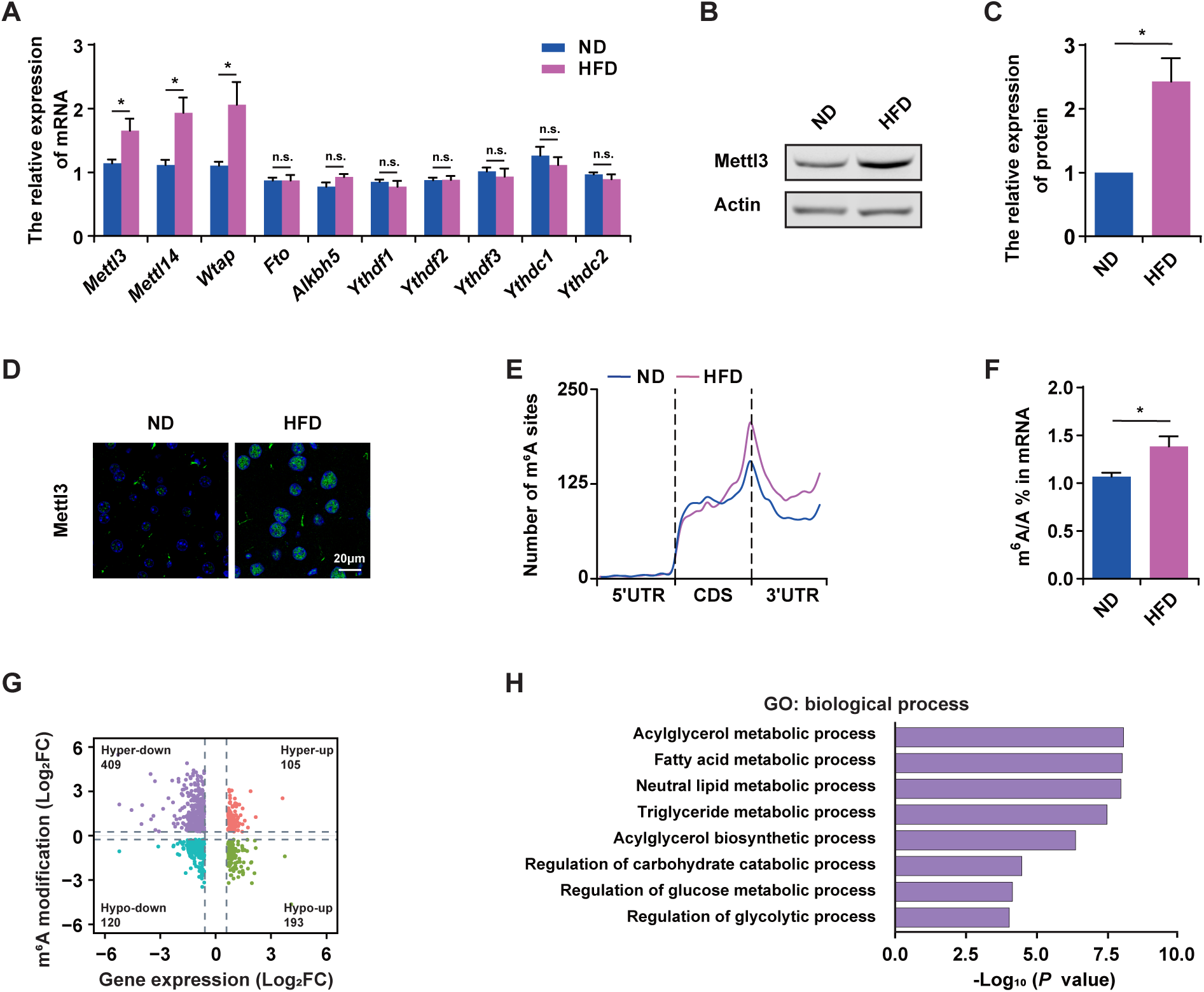
*Mettl3* expression and m^6^A level increased in HFD mice. **A.** qRT-PCR analysis of the expression of *Mettl3*, *Mettl14*, *Wtap*, *Fto*, *Alkbh5*, *Ythdf1*, *Ythdf2*, *Ythdf3*, *Ythdc1*, and *Ythdc2* in the livers of ND and HFD mice, *Ubc* served as the internal control, *n* = 5. **B and C.** Western blotting detection **(B)** and quantification of Mettl3 protein expression **(C)** in the livers of ND and HFD mice, Actin was used as loading control, *n* = 3. **D.** Immunostaining of Mettl3 (green) in the livers of ND and HFD mice. Scale bar, 20 m. **E.** Distribution of m^6^A sites along the 5’UTR, CDS, and μ 3’UTR regions of mRNAs from the ND and HFD mouse livers. **F.** UPLC-MRM-MS/MS showing the percentage of mRNA m^6^A levels in ND and HFD mouse livers, *n* = 4. **G.** Distribution of genes with significant changes in both m^6^A level and gene expression level in HFD condition. **H.** Significantly enriched (*P* < 0.05) gene ontology biological process categories of genes with down-regulated expression and up-regulated m^6^A level in the HFD mouse livers. HFD mice were fed with 60 kcal% fat diet for 20 weeks. Data are presented as mean ± SEM. Unpaired *t*-test, \**P* < 0.05, n.s, no significance. ND, normal diet; HFD, high fat diet; UPLC-MRM-MS/MS, ultra performance liquid chromatography-triple quadrupole mass spectrometry coupled with multiple-reaction monitoring; m^6^A, *N^6^*-methyladenosine; UTR, untranslated regions; CDS, coding sequence; GO, gene ontology. Raw data were displayed in Table S2.

To further investigate the underlying mechanisms of Mettl3 in HFD-induced metabolic disorder, we performed RNA sequencing (RNA-seq) and m^6^A individual-nucleotide-resolution cross-linking and immunoprecipitation sequencing (miCLIP-seq) using mRNAs extracted from the livers of Normal Diet (ND) and HFD mice (Figure S1A). Consistent with previous reports [6, 27], the m^6^A sites in liver mRNAs were also enriched in the regions with RRACH motif (Figure S1B) and tended to occur near stop codons and within 3’UTRs of mRNAs (Figure S1C). More importantly, we detected increased m^6^A sites in the HFD mouse liver **(Figure 1E).** To further validate the presence of m^6^A modifications in the mRNAs of HFD mouse liver, we also applied ultra performance liquid chromatography-triple quadrupole mass spectrometry coupled with multiple-reaction monitoring (UPLC-MRM-MS/MS) analysis to quantify the m^6^A content in mRNAs and observed increased mRNA m^6^A modifications in the HFD mouse liver **(Figure 1F)**, which is consistent with the high expression of Mettl3. 16,686 m^6^A sites were newly induced in the liver mRNAs of HFD mice (Table S1), corresponding to 1860 methylated genes in the HFD mouse liver (Figure S1D). The proportion of unique m^6^A sites and overlaid sites with higher m^6^A level in HFD compared with that in ND mouse liver mRNAs also confirmed the increased m^6^A sites in the HFD mouse liver mRNAs (Figure S1E). To investigate the association of m^6^A with gene expression, we analysed the RNA-seq data for the ND and HFD liver samples, and identified 1913 differentially expressed mRNAs in total with 714 up-regulated genes and 1199 down-regulated genes (rpkm > 1). Meanwhile, we combined the gene expression with m^6^A levels, and 514 genes with increased m^6^A levels in HFD mouse liver were being discovered. Since it has been reported that the presence of m^6^A sites facilitates mRNA degradation [6], we mainly focused on the 409 genes with both hyper m^6^A level and decreased expression in the HFD mouse liver **(Figure 1G)**, as this group of transcripts was likely stabilized after m^6^A depletion. Gene ontology analysis revealed that most of these genes were enriched in lipid metabolic processes, including acylglycerol metabolic process and fatty acid metabolic process, *etc*. Glycometabolism related pathways such as regulation of carbohydrate process were also enriched **(Figure 1H)**. Taken together, m^6^A level and its methyltransferase Mettl3 were consistently up-regulated in the livers of HFD mice, indicating that Mettl3 mediated m^6^A methylation might be involved in metabolic disorders induced by HFD.

### Overexpression of *Mettl3* aggravates liver metabolic disorders and hepatogenous diabetes

To confirm the relationship between high expression level of Mettl3 and HFD-induced metabolic disorders, we specifically overexpressed *Mettl3* in liver by AAV8 which specifically targets hepatocytes [28] and hepatocyte-specific promoter: LP1 [29] (Figure S2A). Living imaging reconfirmed the specifically expressed luciferase in mouse livers at 4 weeks after AAV retro orbital injection, and demonstrated that Mettl3 also specifically expressed in liver (Figure S2B). Moreover, qRT-PCR and western blotting revealed the successfully overexpression of Mettl3 in liver (Figure S2C-E).

We tracked the changes in mouse body weight and metabolic parameters in response to HFD. Compared with mutant Mettl3 conditional overexpression mice (cOE-Mut, served as Ctrl), Mettl3 conditional overexpression mice (cOE) during HFD showed more increase in body weight **(Figure2A)** due to more subcutaneous fat (Figure S2F). The ratio of liver weight to body weight and Oil Red O (ORO) staining further revealed that cOE mice presented more HFD-induced hepatic steatosis **(Figure 2B-D)**. Moreover, compared with cOE-Mut mice, total cholesterol (TC) in serum of cOE mice also increased, while there was no significant change in total triglyceride (TG) **(Figure 2E, F)**.

**Figure 2.**
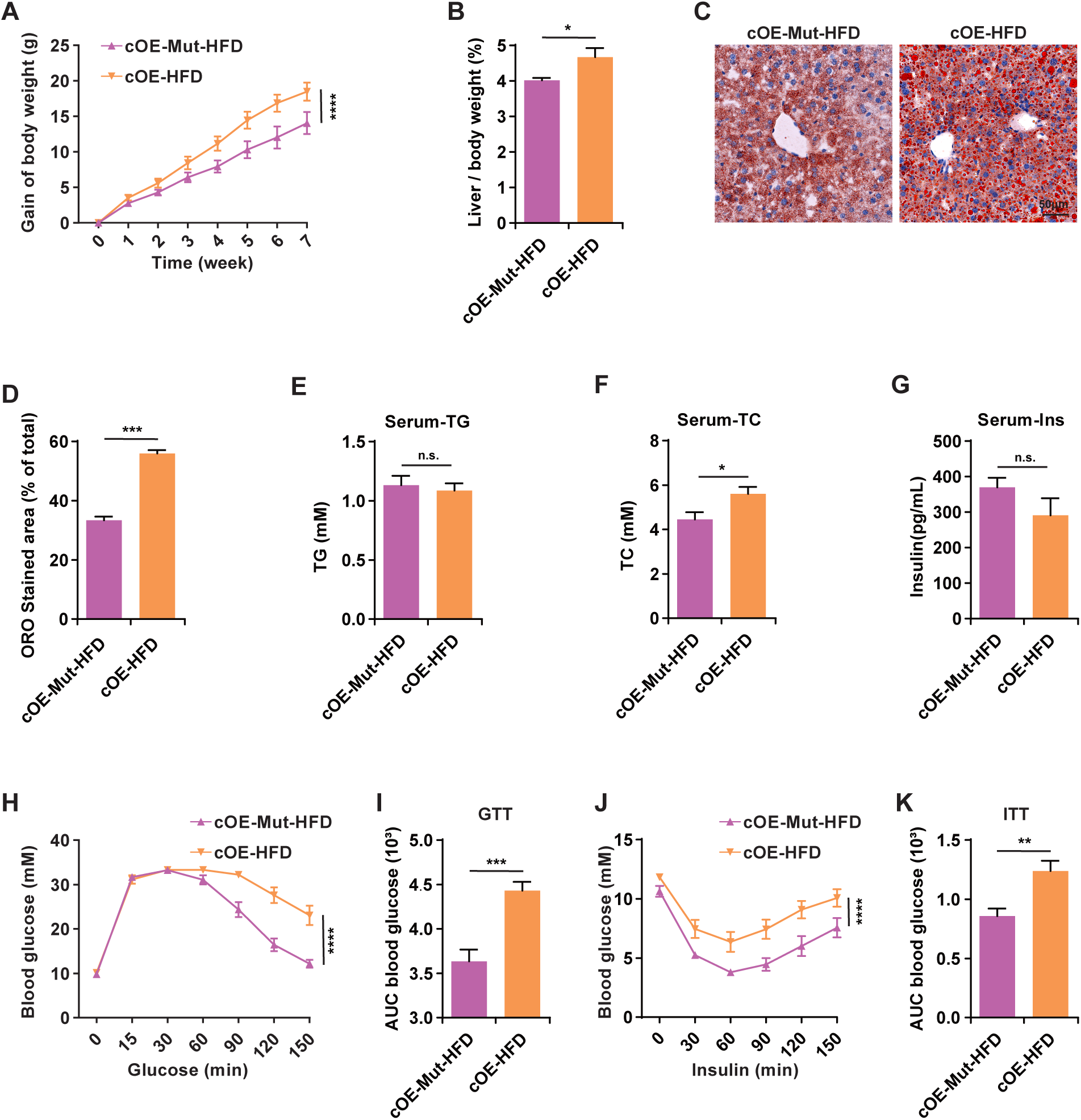
Overexpression of *Mettl3* aggravates liver metabolic disorders and hepatogenous diabetes. **A.** Body weight gain of cOE-Mut and cOE mice during 7 weeks of HFD, *n* = 10. Gain of body weight (g) = final body weight (g) –initial body weight (g) **B.** The ratio of liver weight to body weight of cOE-Mut and cOE mice after 7 weeks of HFD, *n* = 5. **C.** Representative photomicrographs of ORO stained livers of cOE-Mut and cOE mice after 7 weeks of HFD. Scale bar, 50 μm. **D.** Proportion of ORO-stained area in μ cOE-Mut and cOE mouse livers after 7 weeks of HFD, *n* = 3. **E and F**. TG (**E**) and TC (**F**) contents in serum of cOE-Mut and cOE mice after 7 weeks of HFD, *n* = 10. Insulin contents in serum of cOE-Mut and cOE mice after 8 weeks of HFD, *n* = 10. Blood glucose of cOE-Mut and cOE mice after 7 weeks of HFD during GTT, *n* = 8. **I.** AUC statistics for **(H)**, *n* = 8. **J.** Blood glucose of cOE-Mut and cOE mice after 8 weeks of HFD during ITT. *n* = 8. **K.** AUC statistics for **(J)**, *n* = 8. Data are presented as mean ± SEM. Unpaired *t*-test, \**P* < 0.05, \*\**P* < 0.01, \*\*\**P* < 0.001, \*\*\*\**P* < 0.0001. n.s, no significance. cOE-Mut, mutant *Mettl3* (DPPW→APPA) conditional overexpressed mice, served as Ctrl; cOE, *Mettl3* conditional overexpressed mice; ORO, Oil Red O; TG, total triglyceride; TC, total cholesterol; Ins, insulin; GTT, glucose tolerate test; ITT, insulin tolerate test. AUC, area under the curve. Raw data were displayed in Table S2.

Although, there was no significant change in serum insulin level **(Figure 2G)**, further analysis showed that cOE mice presented significant worse glucose tolerance **(Figure 2H, I)** and insulin sensitivity **(Figure 2J, K)** than cOE-Mut mice in HFD condition, evaluated by the glucose tolerance test (GTT) and the insulin tolerance test (ITT), respectively. Together, these results indicated that Mettl3 overexpression can aggravate liver metabolic disorders and hepatogenous diabetes, suggesting that high levels of Mettl3 may be a risk factor for HFD-induced metabolic syndrome.

### *Mettl3* ablation protects mice against HFD-induced metabolic syndrome

Considering that overexpression of Mettl3 can aggravate liver metabolic disorders and hepatogenous diabetes induced by HFD, we supposed *Mettl3* ablation in liver could resistant HFD-induced metabolic syndrome. To verify this hypothesis, we generated *Mettl3* conditional knockout mice (cKO) by crossing *Alb-Cre* and *Mettl3*^flox/flox^ mice (Figure S3A). CRE enzyme specifically expressed in liver and produced *Mettl3* transcripts without exon 2-4, and CRE enzyme didn’t leak into other tissues (Figure S3B). qRT-PCR, western blotting, and immunohistochemistry assays confirmed Mettl3 successfully deletion in liver at both mRNA and protein levels (Figure S3C-F).

As expected, the body weight of cKO mice increased more slowly than Ctrl **(Figure 3A)**, and they also had less subcutaneous fat than Ctrl after HFD for 20 weeks (Figure S3G). HFD-induced hepatic steatosis was slighter in cKO mouse liver, evaluated by the ratio of liver weight to body weight and ORO staining **(Figure 3B-D)**. However, in the late stages of HFD, Mettl3 depletion seemed could not confront the lipid accumulation significantly. We speculate that in the late stages of HFD, lipid accumulation may have reached the limit of liver. However, the damage in Ctrl-HFD mouse liver was severer by showing more and bigger vacuoles **(Figure 3E)**. In addition, although there was no significant change in serum TG, the serum TC of cKO-HFD mice decreased, consistent with the phenotype of cOE mice **(Figure 3F, G)**.

**Figure 3.**
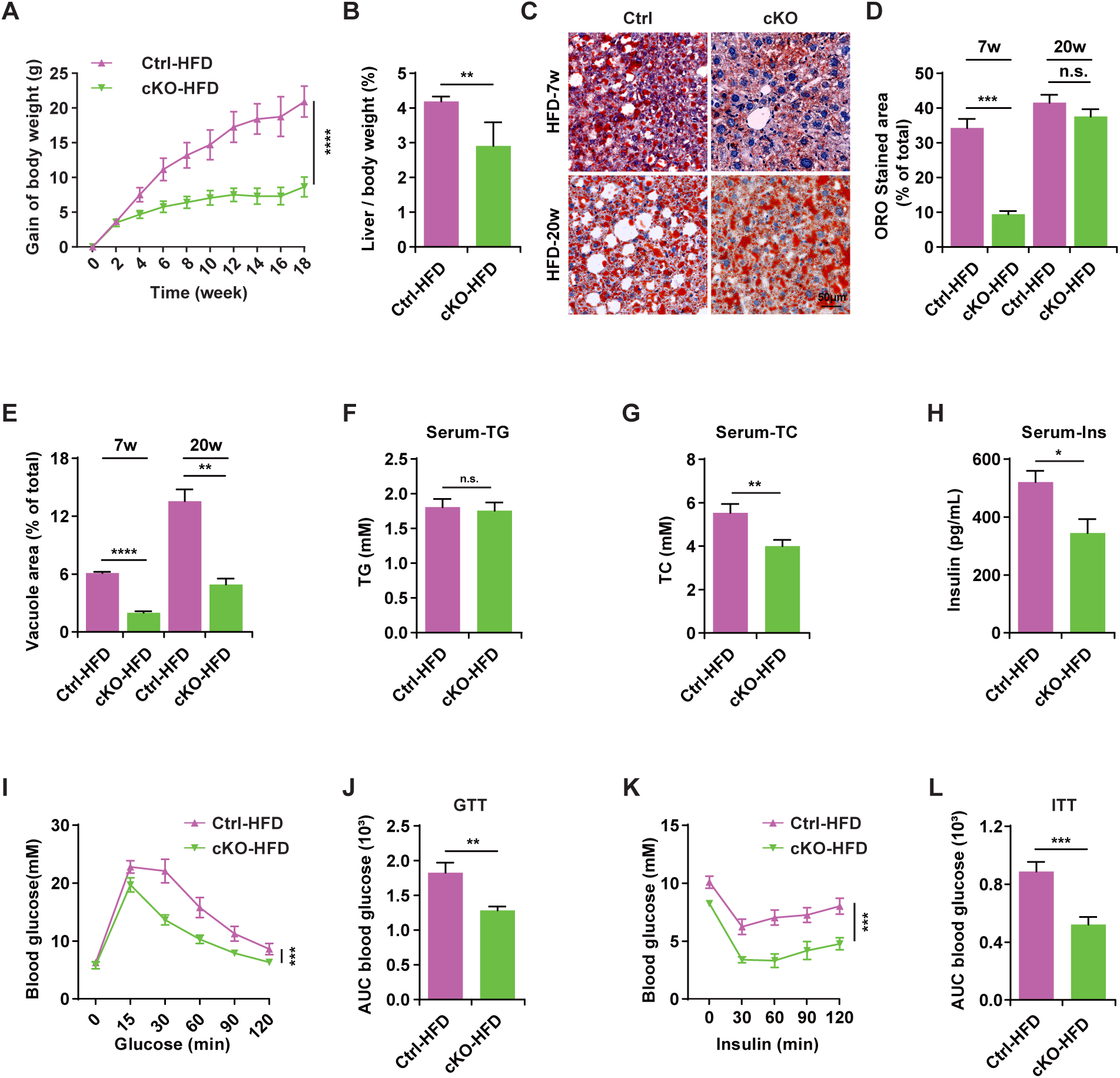
*Mettl3* ablation protects mice against diet-induced metabolic syndrome. **A.** Body weight gain of Ctrl and cKO mice during 20 weeks of HFD, *n* = 10. **B.** The ratio of liver weight to body weight of Ctrl and cKO mice after 20 weeks of HFD, *n* = 5. **C.** Representative photomicrographs of ORO stained livers of Ctrl and cKO mice after 7 weeks and 20 weeks of HFD. Scale bar, 50 m. **D.** Proportion of ORO-stained μ area in Ctrl and cKO mouse livers after 7 weeks and 20 weeks of HFD, *n* = 3. **E.** Proportion of vacuole area in Ctrl and cKO mouse livers after 7 weeks and 20 weeks of HFD, *n* = 3. **F and G.** TG (**F**) and TC (**G**) contents in serum of Ctrl and cKO mice after 20 weeks of HFD, *n* = 8. **H.** Insulin contents in serum of Ctrl and cKO mice after 20 weeks of HFD, *n* = 8. **I.** Blood glucose of Ctrl and cKO mice after 20 weeks of HFD during GTT, *n* = 10. **J.** AUC statistics for **(I)**, *n* = 10. **K.** Blood glucose of Ctrl and cKO mice after 21w of HFD during ITT, *n* = 10. **L.** AUC statistics for (**K**), *n* = Data are presented as mean ± SEM. Unpaired *t*-test, \**P* < 0.05, \*\**P* < 0.01, \*\*\**P* < 0.001, \*\*\*\**P* < 0.0001, n.s, no significance. Ctrl, *Mettl3^flox/flox^* mice; cKO, *Mettl3^flox/flox^; Alb-Cre* mice. Raw data were displayed in Table S2.

It’s worth noting that the serum insulin level significantly decreased in cKO-HFD mice **(Figure 3H)**. Meanwhile, consistent with the glucometabolic phenotype of cOE mice, cKO mice presented significantly better glucose tolerance **(Figure 3I, J)** and insulin sensitivity **(Figure 3K, L)** than Ctrl mice in HFD condition. Taken together, these results suggest that Mettl3 depletion in liver can protect mice against HFD-induced metabolic syndrome, indicating that Mettl3 might be a potential therapeutic target for liver metabolic diseases.

### *Mettl3* ablation alters the expression pattern of lipid and glucose metabolic genes

To further explore the underlying mechanisms of Mettl3 depletion protecting liver from metabolic syndrome induced by HFD, we analysed RNA-seq and miCLIP-seq data generated from livers of Ctrl and cKO mice after 20 weeks of HFD (termed as Ctrl-HFD and cKO-HFD, respectively). Similarly, the m^6^A sites in cKO-HFD mouse liver mRNAs were enriched in the regions with RRACH motif (Figure S4A) and tended to occur near stop codons and within 3’UTRs of mRNAs (Figure S4B). Within all the methylated mRNAs, around 35.1% methylated mRNAs were found to contain one m^6^A site (Figure S4C and Table S1). Since HFD-induced Mettl3 up-regulation and *Mettl3* knockout genetic manipulation have opposite effects on m^6^A level, there was no obvious differences in the number of m^6^A sites and methylated genes between Ctrl-HFD and cKO-HFD mouse livers **(Figure 4A)**. Meanwhile, the m^6^A sites across the entire gene bodies of Ctrl-HFD and cKO-HFD mouse livers also displayed similar distribution **(Figure 4B)**. Meanwhile, it seems that HFD-induced Mettl3 up-regulation played a more dominant role according to the proportion of unique m^6^A sites and overlaid sites with higher m^6^A level in Ctrl-HFD and cKO-HFD mouse livers **(Figure 4C)**. Among the hypo-methylated genes, 212 genes were up-regulated while 116 genes were down-regulated in cKO-HFD mouse liver **(Figure 4D)**. Given that m^6^A is mainly reported to play a negative role in mRNA stability regulation, we focused on the m^6^A-containing up-regulated genes in cKO-HFD mice liver, performed gene ontology analysis, and found that these genes were enriched in insulin response and lipid metabolic related processes **(Figure 4E)**.

**Figure 4.**
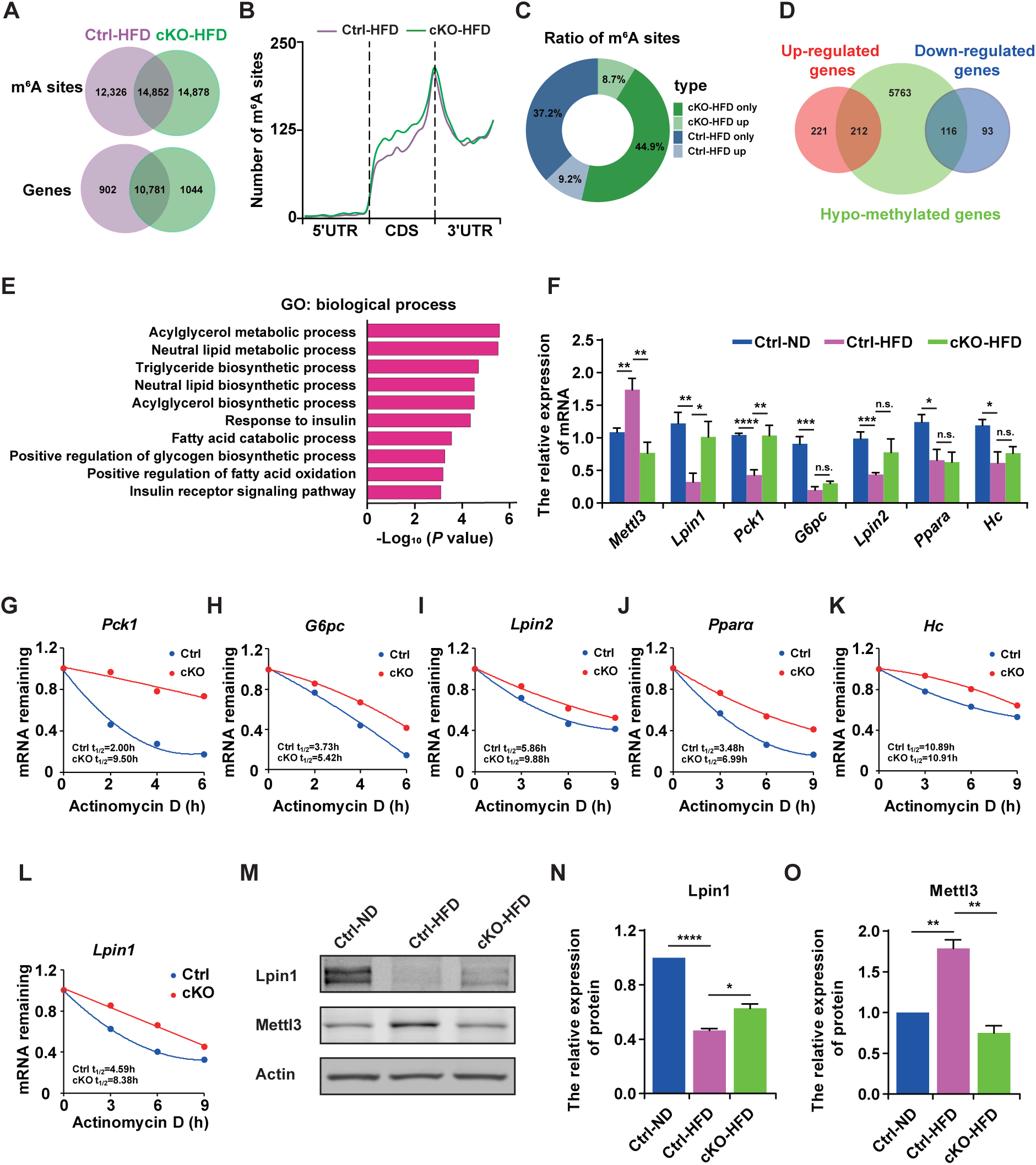
*Mettl3* ablation alters the expression pattern of lipid and glucose metabolic genes. **A.** Venn diagram depicting the overlap of m^6^A sites and methylated genes between liver mRNAs of Ctrl-HFD and cKO-HFD mice. Numbers represent the counts of m^6^A sites or genes in each group. **B.** Distribution of m^6^A sites along the 5’UTR, CDS, and 3’UTR regions of liver mRNAs from Ctrl-HFD and cKO-HFD mice. **C.** Donut charts showing the proportion of unique m^6^A sites and overlaid sites with higher m^6^A level in the livers of Ctrl-HFD and cKO-HFD mice. **D.** Venn diagram representing the relationships between altered genes and m^6^A modification. Numbers represent the counts of genes in each group. **E.** Gene ontology biological process categories (*P* < 0.05) of genes with down-regulated expression and up-regulated m^6^A level in the livers of cKO-HFD mice. **F.** qRT-PCR validation of liver lipid and glucose metabolic genes, with down-regulation in Ctrl-HFD mouse liver (compare with Ctrl-ND) versus up-regulation in cKO-HFD mouse liver (compare with cKO-HFD), *Ubc* served as the internal control, *n* = 5. **G-L.** mRNA half-lives of *Pck1*, *G6pc*, *Lpin2*, *Ppar*, *Hc*, *and* α *Lpin1*. mRNA levels were measured at the indicated time points after Actinomycin D treatment by qRT-PCR, *Ubc* served as the internal control, *n* = 3. **M-O.** Western blotting detection and quantification of Mettl3 and Lpin1 protein expression in liver extracts from Ctrl-ND, Ctrl-HFD, and cKO-HFD mice Actin served as the loading control. *n* = 3. All mice livers were prepared after 20 weeks of HFD. Data are presented as mean ± SEM. Unpaired *t*-test, \**P* < 0.05, \*\**P* < 0.01, \*\*\**P* < 0.001, \*\*\*\**P* < 0.0001, n.s, no significance. Raw data were displayed in Table S2.

qRT-PCR further validated the expression of these candidate genes which were down-regulated in Ctrl-HFD mouse liver (compared with Ctrl-ND) while up-regulated in cKO-HFD mouse liver (compared with Ctrl-HFD), such as *Lpin1*, *Pck1*, *G6pc*, *Lpin2*, *Ppar*, and *Hc* **(Figure 4F)**, and most of these genes were more stable in cKO α mouse liver due to *Mettl3* depletion induced m^6^A loss, indicated by mRNA stability assay **(Figure 4G-L)**. Among them, *Lpin1* has been reported to play an important role in liver lipid metabolism and insulin resistance [30–33], Furthermore, Lpin1 protein also decreased in Ctrl-HFD mouse liver while increased in cKO-HFD mouse liver, contrary to the expression pattern of Mettl3 **(Figure 4M-O)**. Collectively, these findings demonstrated that Mettl3 ablation stabilized key lipid and glucose metabolic genes, especially improved the stability of *Lpin1* mRNA through modulating m^6^A levels.

## Discussion

As the most prevalent mRNA modification in eukaryotes [1], m^6^A involved in many essential biological processes, including cell fate determination [12, 13], embryonic development [13–15] and tumorigenesis [16]. Recent studies have demonstrated that m^6^A modulation of mRNA expression plays an important role in adipogenesis [18], hepatic lipid metabolism [20], obesity [21], and other metabolic diseases, such as NAFLD and type 2 diabetes (T2D) [21–26]. However, the underlying pathways and mechanisms of Mettl3-mediated m^6^A modification regulates liver metabolism remain unclear.

A recent study reported that Mettl3 inhibits hepatic insulin sensitivity via m^6^A located in *Fasn* mRNA and promotes fatty acid metabolism [26]. In our current work, we presented several findings demonstrating the significance of m^6^A in liver metabolic disorders induced by HFD: (1) The major m^6^A methyltransferase Mettl3 and m^6^A level were consistently elevated in the liver from mice after feeding HFD. (2) AAV8 mediated liver conditional overexpression of *Mettl3* aggravated liver and whole-body metabolic disorders, including liver lipid accumulation, abnormal TC in serum and obesity, moreover, *Mettl3* cOE mice presented worse glucose tolerance and insulin sensitivity compared with cOE-Mut mice (served as Ctrl). (3) Mettl3 cKO model generated by crossing *Alb-Cre* and *Mettl3^flox/flox^* mice confirmed that *Mettl3* ablation protects mice against HFD-induced liver metabolic disorders and hepatogenous diabetes. (4) *Mettl3* ablation stabilized key genes involved in liver lipid and glucose metabolism, particularly elevated the stability of an important regulator of liver lipid and glucose metabolism, *Lpin1*. Collectively, our findings demonstrated the critical roles for Mettl3-mediated m^6^A modification in HFD-induced liver metabolic disorders and hepatogenous diabetes, supporting that m^6^A might be a potential therapeutic and diagnostic target for hepatic diseases.

Previous studies have shown that Mettl3 was elevated in peripheral venous blood and livers of T2D patients [26, 34], while the reason for Mettl3 increase was considered as a result of FTO-induced decreases in m^6^A: high-glucose stimulation elevates Fto expression, which leads to m^6^A level decreased, as a response, Mettl3 might increase to maintain the normal m^6^A level [34]. Consistent with these findings, we also detected elevated Mettl3 in livers of mice after feeding HFD. However, the expression of FTO didn’t show significant increase. Therefore, we highly speculate that the up-regulation of Mettl3 in HFD mouse liver may result from other signal pathways.

Several studies showed that Fto-mediated m^6^A demethylation positively regulates adipogenesis, including promoting adipogenesis in porcine intramuscular preadipocytes through inhibiting the Wnt/β-catenin signal pathway [35], increasing adipogenesis in mouse embryonic fibroblasts and primary preadipocytes by regulating mitotic clonal expansion [36], controlling adipogenesis through the regulation of cell cycle in an YTHDF2-m^6^A-dependent manner [37]. Moreover, consistent with these *in vitro* studies, Fto overexpression induced adipocyte hyperplasia in HFD mice [36]. Conversely, Mettl3 negatively correlated with adipogenesis in porcine adipocytes through m^6^A methylation [38], which seems conflict with the weight loss of Mettl3 cKO mice after HFD as we observed. However, obesity is caused by many factors, in this study, it is the result of initial liver metabolic disorders and hepatogenous diabetes, rather than adipogenesis or proliferation of preadipocytes. Meanwhile, the lipid accumulation in hepatocytes is a comprehensive result of liver lipid synthesis, catabiosis, and transportation. Mettl3 is likely to be involved in all these processes and eventually increases lipid accumulation in the HFD mouse liver.

It is interesting to note that lipid accumulation unexpectedly increased in Mettl3 cKO mouse liver in ND condition (data not shown), contrary to the corresponding phenotype of HFD mice. By high throughput RNA-seq and m^6^A-miCLIP-seq, we compared the up-regulated genes with lower m^6^A levels in cKO mouse liver in both ND and HFD conditions, and found that Mettl3 targeted genes were enriched in sterol biosynthetic process in ND condition, but in serval catabolism pathways under HFD condition, such as fatty acid catabolic process and positive regulation of fatty acid oxidation. These findings indicate that Mettl3 might regulate different subsets of genes in different diet conditions and serves as a bidirectional switch in lipid metabolism.

Taken together, we conclude that Mettl3 serves as an essential regulator of liver lipid and glucose metabolism and protects from metabolic disorders and hepatogenous diabetes induced by HFD, thus promoting Mettl3-mediated m^6^A as a target for hepatic diseases’ therapy and diagnosis.

## Materials and methods

### Mouse

The mice used in this study were C57BL/6 strains. Specific pathogen-free-grade mice were purchased from Beijing Charles River Laboratory Animal Center and housed in the animal facilities of the Institute of Zoology, Chinese Academy of Sciences. All animal experiments were carried out under the guidelines for the Use of Animals in Research issued by the Institute of Zoology, Chinese Academy of Sciences.

### Mouse breeding

*Mettl3^flox/+^* mice were generated by the CRISPR-Cas9 system-assisted homologous recombination as previously described [39]. C57BL/6 *Alb-Cre* transgenic mice were purchased from Shanghai BRL Medicine Company. *Mettl3^flox/flox^* mice were obtained by mating *Mettl3*^flox/+^ to each other, *Mettl3*^flox/+^*Alb*-Cre mice were obtained by mating *Mettl3*^flox/flox^ and *Alb*-Cre mice. *Mettl3*^flox/+^ *Alb*-Cre and *Mettl3*^flox/flox^ were crossed to generate *Mettl3*^flox/flox^ *Alb*-Cre mice (Mettl3 cKO).

### Genotyping of mice

All mice were genotyped with the tail DNA which was extracted using the Mouse Direct PCR Kit (Bimake). Briefly, mouse tails were mixed with 50 μL Buffer L and 1 μL Protease Plus, and incubated at 55 °C for 30 min, then 100 °C for 5 min according to the manufacturer’s instructions.

Two pairs of primers were used to detect the loxp insertion into the *Mettl3* intron 1 (L-loxp-F and L-loxp-R) and intron 4 (R-loxp-F and R-loxp-R). The product sizes were 222 bp and 335 bp with the loxp sequence insertion into *Mettl3* intron 1 and intron 4, respectively; whereas the product sizes from WT were 182 bp and 295 bp, respectively. Cre recombinase was detected by the *Alb-Cre* primers and the product of PCR was 350 bp. Heart, liver, spleen, lung, and kidney were extracted to confirm the deletion of *Mettl3* with the primers of L-loxp-F and R-loxp-R and the product of *Mettl3* deletion was 318 bp whereas the WT product was 2554 bp. All primers were listed in Table S3.

### RNA extraction and qRT-PCR

Total RNA was extracted with TRIzol reagent (Invitrogen, 15596-018) from the whole liver and reverse-transcribed into cDNAs using the Reverse Transcription System (Promega, A3500). qRT-PCR was performed using SYBR Premix Ex Taq kit (TaKaRa, RR420A) on Agilent Stratagene Mx3005P. Relative gene expression was analysed based on the 2^−Ct^ method with *Ubc* as the internal control. All primers were listed in Table S3.

### Western blot

Western blotting was performed as described previously [40] with corresponding antibodies: anti-Mettl3 (1:500, Abcam, ab195352), anti-Lpin1 (1:500, Cell Signalling Technology, 5195S), anti-β-Actin (1:2000, Sigma, A1978) and anti-α-Tubulin (1:2000, Sigma,T6199).

### Immunohistochemical analysis

Immunohistochemical was performed as described previously [40]. Anti-Mettl3 (1:500, Abcam, ab195352) antibody and Hoechst 33342 (Invitrogen, H3570) were used. Images were obtained using standard methods with a Leica Aperio VERSA 8 microscope (Leica Biosystems).

### Plasmid construction and virus production

pX602 backbone was modified from pX602-AAV-TBG::NLS-SaCas9-NLS-HA-OLLAS-bGHpA;U6::BsaI-sgRNA, which was a gift from Feng Zhang (Addgene plasmid # 61593; http://n2t.net/addgene:61593; RRID:Addgene_61593) [41], while the LP1 promoter was constructed as previously described [29]. Mettl3 catalytic mutant (395-398 aa, DPPW → APPA) was also generated as previous work [42].

AAV8 was generated with HEK-293 cells, purified with chloroform, and titered by qPCR as previously described [43], then retro orbital injected into mice at the titer of 2.5×10_12_ vg each mouse.

### Oil Red O staining

Liver lipid accumulation was confirmed by Modified Oil Red O stain kit (Solarbio, G1261) according to the manufacturer’s instructions. In brief, frozen slices of liver (6-10 μm) were fixed in 10% formaldehyde in PBS for 10 min, then washed with 60% isopropanol for 20-30 s. Liver tissue was stained in Modified Oil Red O solution for 10-15 min. After staining, the slices were washed with 60% isopropanol and then with H_2_O. Images were obtained using standard methods and imaged with a Leica Aperio VERSA 8 microscope, then analysed with Image J (Media Cybernetics).

### Metabolic measurements

For GTT assay, mice were fasted overnight (for 12 h) and then injected intraperitoneally (i.p.) with D-glucose (2 g/kg body weight). For ITT assay, mice were randomly fed and injected i.p. with insulin from porcine pancreas (0.75 U/kg body 0 weight, Aladdin, I113907). Blood from a tail vein was collected before injection and at different time points after injection (as indicated in the Figures). Glucose concentrations were measured with an AccuCheck blood glucose meter (Roche Diagnostics Inc.). Serum TG and TC concentrations were measured with Automatic biochemical analyzer (Chemray 240). Serum insulin concentrations were measured by the Insulin test ELISA kit (USCN KIT INC, CEA448Mu) and performed as manufacturer’s instructions.

### Fat volume measurement

Mice were anesthetized with isoflurane and put into the Quantum FX system (PerkinElmer, PE Quantum FX) then scanned with X-ray. Data were analysed with Analyze 12.0.

### UPLC-MRM-MS/MS analysis

mRNAs were purified from total RNAs using Dynabeads mRNA purification kit (Ambion, 61006). 200 ng mRNA was mixed with 0.1 U Nuclease P1 (Sigma) and 2.0 U calf intestinal phosphatase (New England Biolabs), in the final reaction volume of 40 μL adjusted with water, and incubated at 37 °C overnight. The mixture was transferred to ultrafiltration tubes (MW cutoff of 3 kDa, Pall, Port Washington, NY) and centrifuged at 4 °C, 14,000 × g for 25 min.

The UPLC-MRM-MS/MS analysis was performed according to a previous report [44]. The LC was performed on an ExionLCTM analytical (UPLC) system (AB Sciex, USA). Chromatographic separation was carried out on an Acquity UPLC HSS T3 column (1.8 μm, 100 mm × 2.1 mm id, Waters, made in Ireland). The flow rate was 0.25 mL/min, and the mobile phase consisted of methanol (solvent A) and water containing 0.1% formic acid (solvent B) in a linear gradient. The gradient program was as follows: 0-2.5 min, 4% A; 2.5-2.7 min, 4 to 31% A; 2.7-6 min, 31% A; 6-6.2 min, 31 to 95% A; 6.2-9.3 min, 95% A; 9.3-9.6 min, 95 to 4% A; 9.6-14.5 min, 4% A. The column temperature was maintained at 40 °C. The temperature of the autosampler was set at 4 °C, and the injection volume was 4 μL.

MS/MS analysis was carried out on a Qtrap 4500 mass spectrometer (AB Sciex, USA) equipped with Turbo Ionspray interface operating in positive ESI mode. The instrument was operated with an ion spray voltage of 4.5 kV and a heater gas temperature of 500 °C. A nebulizer gas (gas 1) of 40 psi, a heater gas (gas 2) of 50 psi, a curtain gas of 20 psi, and a medium collision gas were used. Mass-dependent parameters such as the declustering potential, entrance potential, collision energy, and collision cell exit potential, were set to the optimal values obtained by automated optimization. A multiple reaction monitoring (MRM) mode was employed for data^6^ 136.1 for acquisition. m/z 282.1→150.1 for m A (collision energy, 12 eV), m/z 268.1→A (9 eV). The injection volume for each sample was 5 μL, and the amount of m^6^A μ and A was calibrated by standards curves. The dwell time for each transition was 100 ms. Data acquisition was performed with Analyst 1.6.2 software (Applied Biosystems, USA).

### mRNA stability assay

Primary hepatocytes were plated on 6-well plates with 5 × 10^5^ cells per well and cultured for 2 days. Then cells were treated with actinomycin-D (10 μg/mL, Sigma) and collected at the indicated time points (2 h, 4 h, and 6 h or 3 h, 6 h, and 9 h). Total RNA was extracted and analysed by qRT-PCR. *Ubc* was used as an internal control.

The half-life was calculated as previously described [40]. Three replicates were conducted for each calculation.

### RNA-seq and m^6^A-miCLIP-seq

RNA-seq libraries were directly generated using the KAPA Stranded mRNA-Seq Kit (KAPA, KK8401) following the manufacturer’s instructions.

The preparation of m^6^A-miCLIP-seq libraries were carried out following previously reported methods [45, 46] with some modifications. Briefly, mRNAs purified using Dynabeads mRNA Purification Kit (Life Technologies, 61006) were fragmented to a size of around 100 nt with the fragmentation reagent (Life Technologies, AM8740). 2 μg of purified mRNAs were mixed with 5 μg of anti-m A antibody (Abcam, ab151230) in 450 μL immunoprecipitation buffer (50 mM Tris, pH 7.4, 100 mM NaCl, 0.05% NP-40) and incubated by rotating at 4 °C for 2 h. The solution was then transferred to a clear flat-bottom 96-well plate (Corning) on ice and irradiated three times with 0.15 J/cm^2^ at 254 nm in a CL-1000 Ultraviolet Crosslinker (UVP). The mixture was then immunoprecipitated through incubation with Dynabeads Protein A (Life Technologies, 1001D) at 4 °C for 2 h. After extensive washing and on-bead end-repair and linker ligation, the bound RNA fragments were eluted from the beads by proteinase K digestion at 55 °C for 1 h. RNAs were isolated by further phenol-chloroform extraction and ethanol precipitation. Purified RNAs were used to construct the library using SMARTer smRNA-Seq Kit for Illumina (Takara, 635029) according to the manufacturer’s instructions. Sequencing was carried out on Illumina HiSeq X-ten platform with paired-end 150 bp read length.

### Analysis of RNA-seq data

All the RNA-seq samples were sequenced by Illumine Hiseq 2000 with paired end 150-bp read length. Clean fastq reads after quality control by cutadapt and trimmomatic[48] were aligned to mice reference genome (GRCm38/mm10; Ensemble version 68) via hisat2 (v2.0.5) [47] aligner with default settings. Only the reads with mapping quality score (MAPQ) ≥ 20 were kept for the downstream analysis.

FeatureCounts (v1.6.0) [49] was employed to estimate the read counts of per gene according to library type. Differentially expressed genes was identified by edgeR (v3.18.1) [50] with fold change > 1.5 and *P* value < 0.05 as thresholds between HFD WT and HFD cKO group. In the whole process, we only kept the genes with reads per kilobase per million mapped reads (RPKM) > 1 as the candidate genes for further analysis. Gene ontology (biological process) enrichment analysis (*P* < 0.05) was performed using the R package ClusterProfiler [51].

### Analysis of miCLIP-seq data

#### Read processing

Raw sequencing data quality control was performed by FASTQC. Adaptors were trimmed by fastx_clipper tool from FASTX-Toolkit (http://hannonlab.cshl.edu/fastx_toolkit). For the forward reads, PCR-amplified reads were removed by fastq2collapse.pl from CTK Tool Kit (v1.0.3) [52] via barcode sequence. Cutadapt (v1.16) [48] was employed to trim the polyA-tail. Reverse reads were reversely complemented by fastx_reverse_complement tool from fastx_toolkit and processed in the same way. Random barcode removal was accomplished by stripBarcode.pl from CTK Tool Kit (v1.0.3), only reads longer than 18 nt were kept by Trimmomatic (v0.33) [53].

#### Mapping and mutation calling

Replicate samples were merged and aligned to mice reference genome (GRCm38/mm10; Ensemble version 68) by BWA (v0.7.17-r1188) [54] with the recommend parameter: -n 0.06 -q 20. Cross-linking-induced mutation sites (CIMS) were detected by the CTK Tool Kit (v1.0.3) [52] as reported. For each detected mutation site, the CIMS software identifies the coverages of unique tags (k) and mutation position (m), we only kept the sites with an m/k ratio 1-50% and mutation sites within the RRACH motif as reliable m^6^A sites for subsequent analysis in order to reduce false positives rates [55]. m^6^A sites annotation was performed by intersectBed from BEDTools (version 2.16.2) [56]. The m^6^A motif was generated by WebLogo3 [57]. For the differential m^6^A methylation sites, we obtained the overlaid m^6^A sites between control and condition samples, the read counts span per m^6^A site were calculated by the bedtools multicov tool (version 2.16.2) [56] from miCLIP-seq and RNA-seq uniq reads and divided by the library size, m^6^A enrichment values difference for each site was determined by chi-square test with *P* < 0.05 and 1.2-foldchange as thresholds.

### Statistical analysis

All data are expressed as mean ± SEM. GraphPad Prism 8 (GraphPad Software Inc.) was used for statistical analysis. Unpaired student’s t-test was used to determine the differences between two groups; a two-way ANOVA analysis followed by Bonferroni multiple-comparison test was used to determine differences between multiple groups. *P* < 0.05 was considered statistically significant.

### Data availability

The accession number for the RNA-seq and miCLIP-seq data in this paper is GSA: CRA002000. These data have been deposited in the Genome Sequence Archive under project PRJCA001786.

## Authors’ contributions

Li Y performed the molecular genetic studies, measured mouse metabolic phenotype, participated in the design of the study and drafted the manuscript. Zhang Q performed bioinformatics analysis. Cui G performed m^6^A-miCLIP-seq assay. Zhao F and Tian X performed the UPLC-MRM-MS/MS analysis. Sun BF supported the bioinformatics analysis. Li W and Yang Y conceived the project, supervised the study, interpreted the data, wrote and provided final approval of the manuscript. All authors read and approved the final manuscript.

## Acknowledgements

This work was supported by the Strategic Priority Research Program of the Chinese Academy of Sciences (XDA16030000); the National Key Research and Development Program (2017YFA0103803, 2018YFA0107703, 2018YFA0801200); the National Natural Science Foundation of China (31621004, 31770872); the Key Research Projects of the Frontier Science of the Chinese Academy of Sciences (QYZDY-SSW-SMC002, QYZDB-SSW-SMC022), and the Youth Innovation Promotion Association of Chinese Academy of Sciences (CAS2018133). We thank Dr. Kai Xu from Institute of Zoology, CAS for offering *Mettl3^flox/flox^* mice.

## Supplementary materials

**Figure S1.**
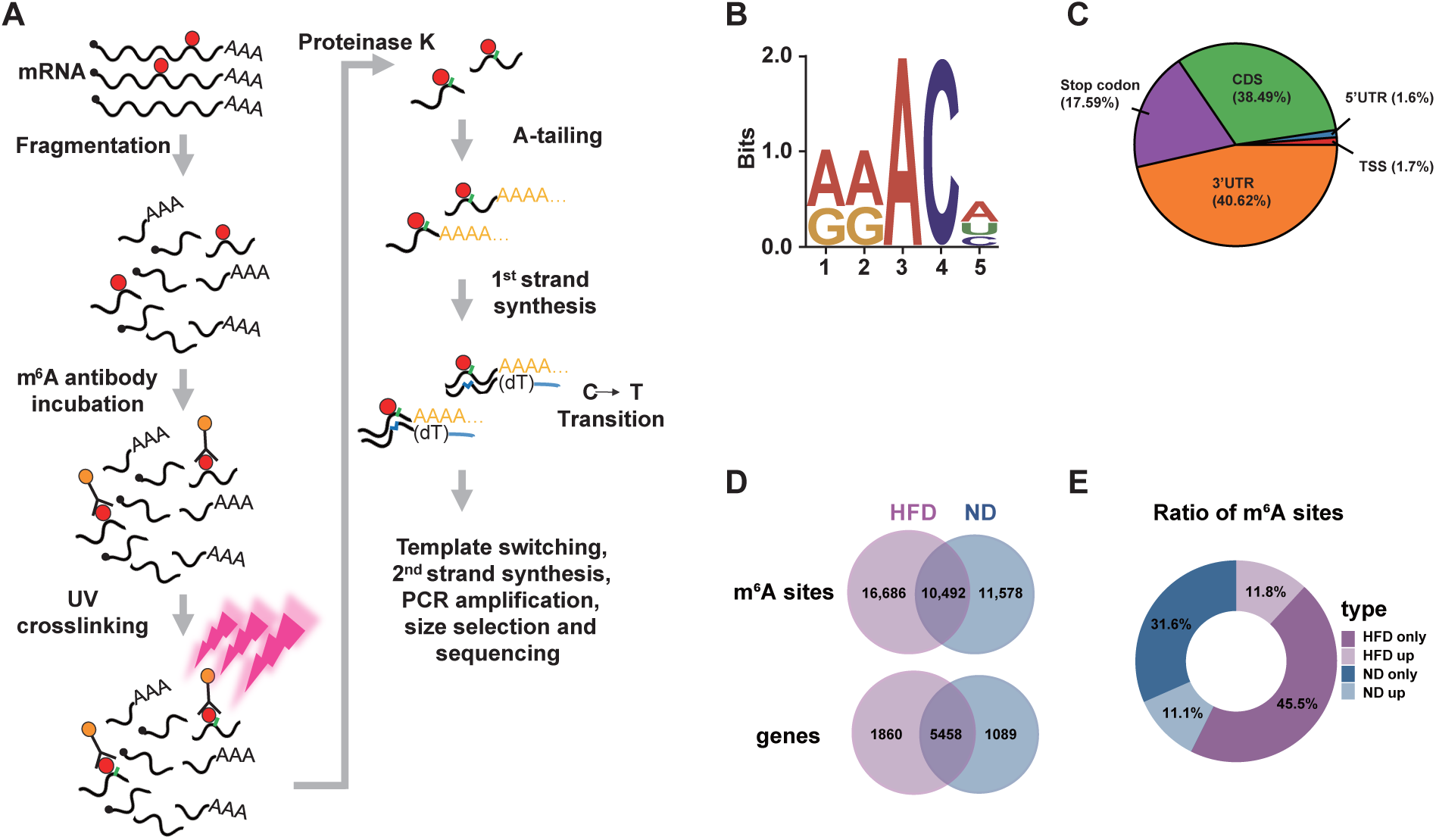
m^6^A pattern in HFD mouse liver. **A.** Schematic illustration of miCLIP library construction procedure. **B.** m^6^A consensus motif in mRNAs from HFD mouse livers. **C.** Transcriptome-wide distribution of m^6^A sites. Pie chart showing the proportion of m^6^A sites in distinct non-overlapping segments: 5’UTR, TSS, CDS, Stop codon and 3’UTR. **D.** Venn diagram depicting the intersection of m^6^A sites and methylated genes between liver mRNAs of ND and HFD mice. Numbers represent the counts of m^6^A sites or genes in each group. **E.** Donut charts showing the ratio of unique m^6^A sites and overlaid sites with higher m^6^A level in the livers of ND and HFD mice. TSS, transcriptional start site.

**Figure S2.**
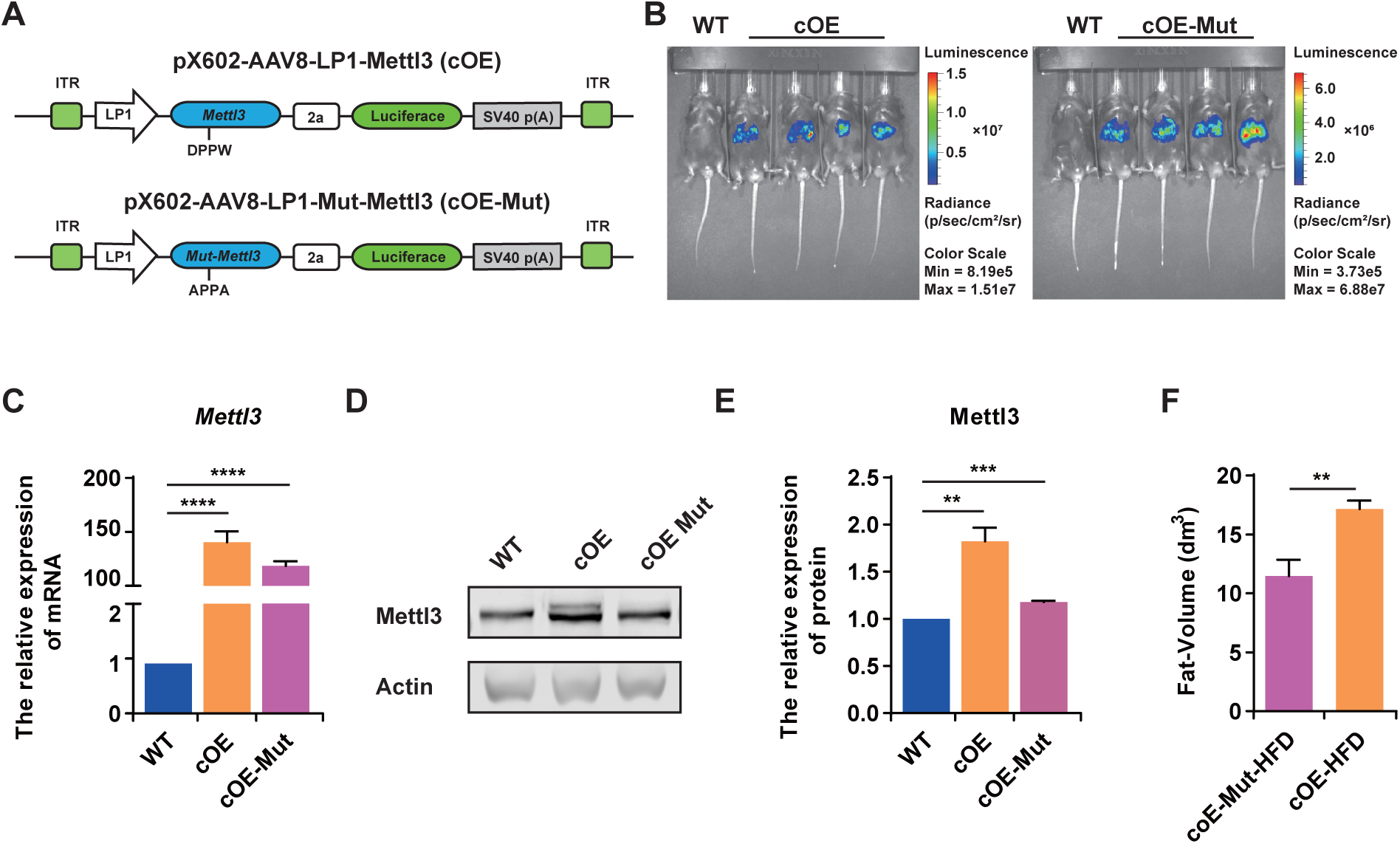
Identification and metabolic indexes of *Mettl3* cOE mice. **A.** Schematics of vectors for conditional overexpression of *Mettl3* (pX602-AAV8-LP1-*Mettl3*, cOE) or Mut-*Mettl3* (pX602-AAV8-LP1-Mut-*Mettl3*, cOE-Mut). cOE-Mut mice served as the control for the following experiments. **B.** Living imaging demonstrating luciferase specifically expressed in cOE and cOE-Mut mouse livers. **C.** qRT-PCR validation of *Mettl3* and Mut-*Mettl3* conditional overexpression in cOE and cOE-Mut mouse livers, *n* = 3. **D and E.** Western blotting detection and quantification of the expression of Mettl3 and mutant Mettl3 proteins in liver extracts from wild type (WT), cOE and cOE-Mut mice. Actin served as the loading control, *n* = 3. **F.** Fat-Volume of cOE-Mut and cOE mice after 7 weeks of HFD, *n* = 10. Data are presented as mean ± SEM. Unpaired *t*-test, \*\**P* < 0.01, \*\*\**P* < 0.001, \*\*\*\**P* < 0.0001, n.s, no significance. ITR, inverted terminal repeats. Raw data were displayed in Table S2.

**Figure S3.**
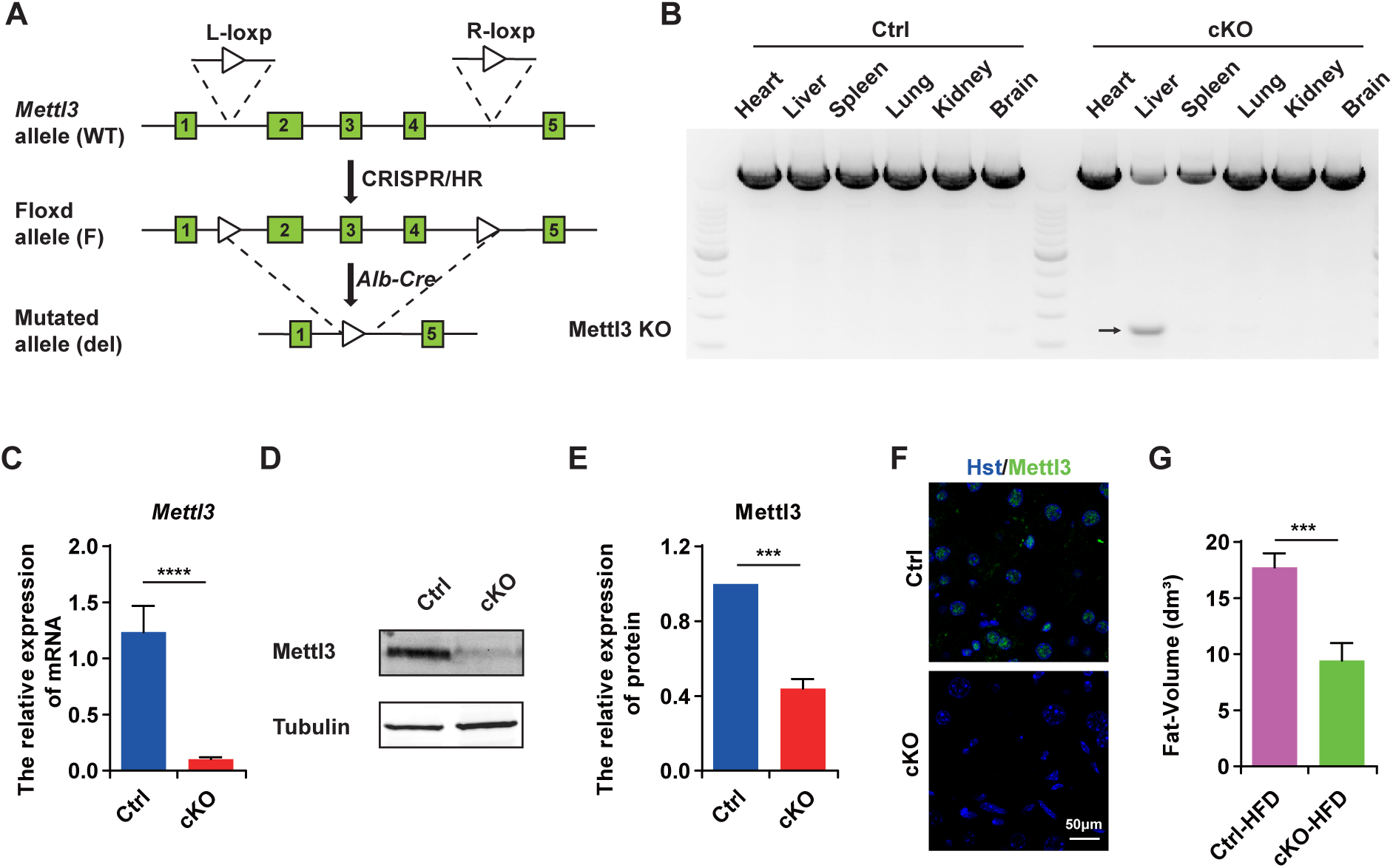
Identification and metabolic indexes of Mettl3 cKO mice. **A.** Diagram illustrating the procedure of generating *Alb-Cre* mediated *Mettl3* conditional knockout mice. **B.** PCR confirmed the conditional knockout of *Mettl3* exons 2-4 in the livers of *Mettl3^flox/flox^; Alb-Cre* (cKO) mice. **C.** qRT-PCR showing the down-regulated of *Mettl3* mRNA in cKO mouse livers, *n* = 3. **D and E.** Western blotting detection and quantification of Mettl3 protein expression in the liver extracts from Ctrl and cKO mice. Tubulin served as the loading control. *n* = 3. **F.** Immunostaining of Mettl3 (Green) from Ctrl and cKO mouse livers. Scale bar, 50 μm. **G.** Fat-Volume of Ctrl and cKO mice after 20 weeks of HFD, *n* = 10. Data are presented as mean ± SEM. Unpaired *t*-test, \**P* < 0.05, \*\*\**P* < 0.001, n.s, no significance. Raw data were displayed in Table S2.

**Figure S4.**
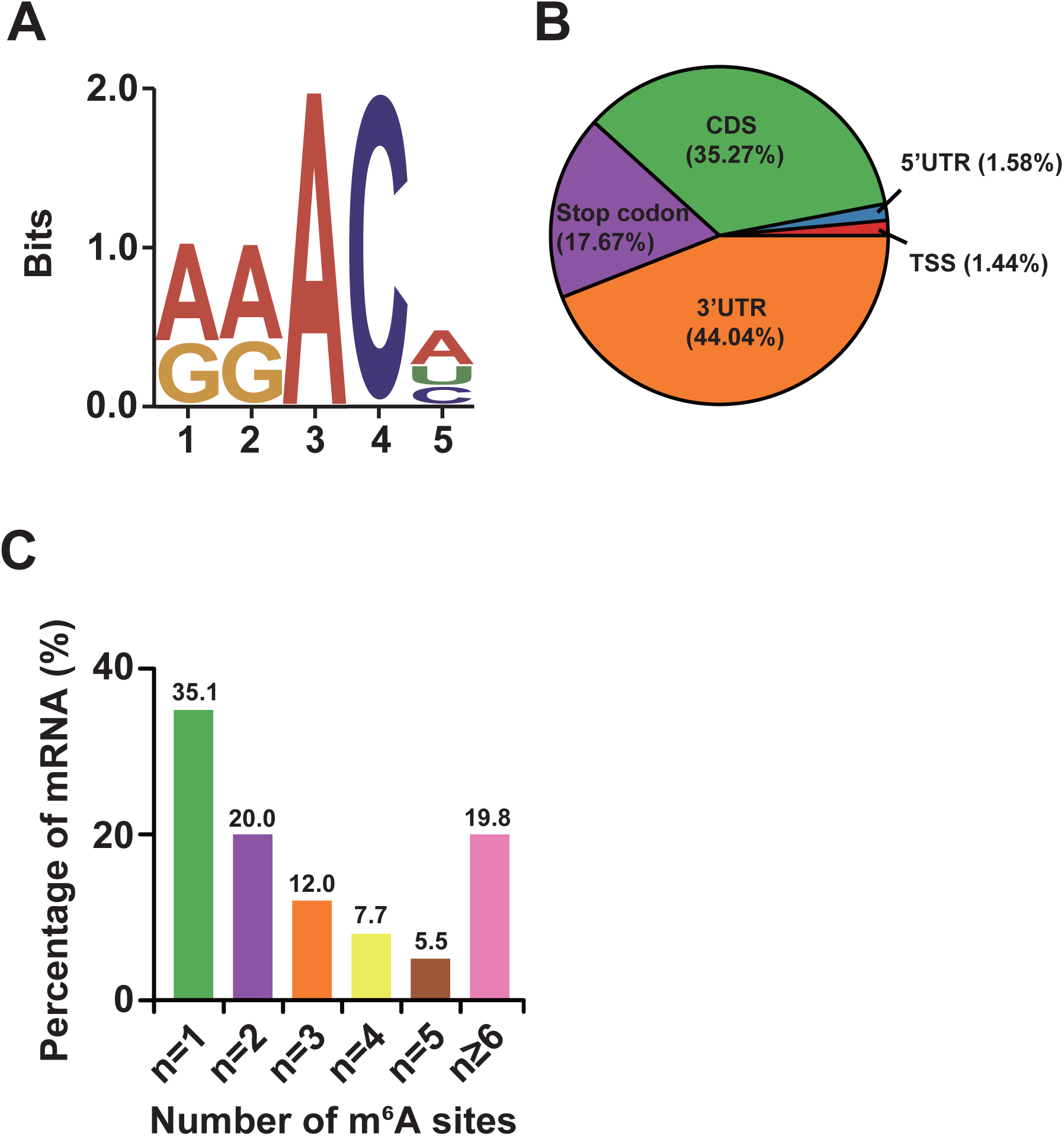
m^6^A pattern in liver of *Mettl3* cKO mouse after HFD. **A.** m^6^A consensus motif in mRNAs from liver of Mettl3 cKO mouse after 20 weeks HFD. **B.** Transcriptome-wide distribution of m^6^A sites of mRNAs from liver of Mettl3 cKO mouse after 20 weeks HFD. Pie chart showing the proportion of m^6^A sites in distinct non-overlapping segments: 5’UTR, TSS, CDS, Stop codon and 3’UTR. **C.** Number of m^6^A-methylated mRNAs with different numbers of m^6^A sites.

## Supplementary Tables

**Table S1 The identified m^6^A sites in ND, HFD and cKO-HFD mice.**

**Table S2 Statistics source data.**

**Table S3 The information of primers used in this study.**

